# A Machine Learning Approach to Predict Cellular Mechanical Stresses in Response to Chemical Perturbation

**DOI:** 10.1101/2023.02.02.526750

**Authors:** V.A. SubramanianBalachandar, Md. Mydul Islam, R. L. Steward

## Abstract

Mechanical stresses generated at the cell-cell level and cell-substrate level have been suggested to be important in a host of physiological and pathological processes. However, the influence various chemical compounds have on the mechanical stresses mentioned above is poorly understood, hindering the discovery of novel therapeutics and representing a barrier in the field. To overcome this barrier, we implemented two machine learning (ML) Models: Stepwise Linear Regression (SLR) and Quadratic Support Vector Machine (QSVM) to predict the dose-dependent response of tractions and intercellular stresses to chemical perturbation. We used traction and intercellular stress experimental data gathered from 0.2 μg/mL and 2 μg/mL drug concentrations along with cell morphological properties to train our model. To demonstrate the predictive capability of our ML model we predicted tractions and intercellular stresses in response to 0 μg/ml & 1 μg/ml drug concentrations. Results revealed the QSVM model to best predict intercellular stresses, while SLR best predicted tractions.

**Author Summary:** The ML framework we present here can be used to predict the mechanical response of any anchorage-dependent cell type to any chemical perturbation. The proposed ML can directly predict the intercellular stresses or tractions as a function of drug dosage and/or monolayer/cell coverage area which could potentially reduce the experimental time on studying the mechanics of cells to external chemicals or mechanical constraints. We believe our findings could be helpful in accelerating drug discovery and increase our understanding in the role of cellular stresses during disease progression.

## Introduction

The mechanosensing ability of cells is critical for many biological processes such as cell migration, growth, and differentiation and has physiological and pathological implications [1]. As the adherent cell migrates through and probes its environment contractile forces must be generated and when migrating as a collective these same adherent cells also interact with each other by transmitting intercellular stresses through cell-cell junctions. Cell-cell junctions enable fast, long distance mechanical force communication which subsequently yields intercellular stresses [2]. We measure intercellular stresses using monolayer stress microscopy (MSM) and tractions using traction force microscopy [1].

Tractions are generated via actomyosin contractility and actin polymerization and were initially measured by observing wrinkles exerted by single cells on thin, silicone membranes [3]. This method would later be extended to measure tractions generated by cells attached to extracellular matrix (ECM) coated polymers and tractions generated by cells attached to flexible, micropost force sensor arrays [3]. Three dimensional tractions have also been measured by cells culture on top of flexible polymers and cells embedded in 3-D matrices [4]. A novel Förster resonance energy transfer (FRET) sensor-based approach has also been employed to measure tractions by estimating the change in excitation energy of fluorescent protein markers that are sensitive to external forces [5, 6]. The many variations by which tractions can be measured has led to a host of studies revealing the importance of tractions both physiologically and pathologically. In fact, tractions have been shown to be important in cell adhesion, spreading, migration, and ECM remodeling [7–10] and is linked to various pathologies including cancer metastasis, fibrosis, and inflammation [11–14].

For cells in a monolayer in addition to tractions being generated at the cell-substrate interface, at the cell-cell interface intercellular stresses are generated. Intercellular stresses have been suggested to be important in tissue morphogenesis, epithelial–mesenchymal transition, wound healing and tumor progression [15–17]. Intercellular stresses can be calculated using MSM which was first described by Tambe et al [18]. In brief, both normal and shear intercellular stresses are recovered from tractions by assuming a monolayer sheet of cells as elastic thin plates and imposing Newton’s force balance and strain compatibility equations [19]. However, other groups would also develop additional alternative methods to calculate intercellular stresses as well as discussed below.

More recently, Bayesian Inversion Stress Microscopy (BISM) [20] and Kalman Inversion Stress Microscopy (KISM) [21] were presented as predictive models for internal stress field based on corresponding traction force data. BISM and KISM use experimental traction force data as a likelihood function to make intercellular stress field predictions using Bayesian statistics (Bayes’ theorem). BISM can infer internal stress fields only from a single traction field image, but a dimensionless regularization parameter must be calculated from the experimental data to make predictions. KISM however, is capable of estimating internal stress fields from a time lapse of traction data (movie) with its accuracy depending on time resolution of the traction data [20, 21]. There are several mechanical factors notably substrate stiffness, cell area, local cell curvature, and external forces that have demonstrated to affect tractions and intercellular stresses [22–27]. Ghosh et al., predicted endothelial cell tractions by using substrate stiffness and cell area [27]. A positive correlation was observed for both stiffness and area with respect to the tractions [23]. Contrastingly, [24] reported that larger cell area lowered average traction forces in human pulmonary artery ECs.

The physiological relevance of tractions and intercellular stresses are equally as important as their pathological ramifications. For example, upregulation of endothelial contractility and increase in tractions via actin stress fiber and ECM remodeling are linked to higher cellular and vascular stiffness and vascular hyperpermeability as seen in hypertension and atherogenesis [28–31]. Furthermore, tractions and intercellular stresses have been linked to vascular hyperpermeability via ROCK1/2 and thrombin mediated pathway [32]. Such hyperpermeability induces loss of blood brain barrier integrity and has also been linked with several neurological disorders such as multiple sclerosis, stroke, and traumatic brain injury [33].

Taken into account the physiological and pathological relevance of both tractions and intercellular stresses, we propose that TFM and MSM are powerful tools that can be utilized to clarify the biomechanical mechanisms of various diseases, some of which have been mentioned above and potentially lead to novel therapeutics. Thus far, the development of novel regenerative medicine and drugs have been heavily researched to treat numerous vascular-related diseases such as hypertension, atherosclerosis, stroke, and coronary artery disease (CAD), for example [34–36]. However, a barrier exists in the field as each of these mechanics-related diseases often require treatment with drugs, whose influence is dose-dependent. We propose that studying the biomechanical mechanism by which certain drugs influence cells and the subsequent tissues they constitute could improve drug efficacy. However, the time and the financial resources required to evaluate the impact of various drugs on cell behavior can be overwhelming and expensive. Machine learning offers an alternative approach that has the potential to resolve or at the bare minimum mitigate the issues previously mentioned. It was therefore our objective to apply machine learning approach to predict the dose-dependent cellular, biomechanical response to chemical stimulation.

In this paper, we utilized machine learning to predict both tractions and intercellular stresses as a function of drug concentration and cell morphological parameters such as monolayer perimeter and cell area. Predictive models were created using Stepwise Linear Regression (SLR) and Quadratic Support Vector Machine (QSVM) regression learners. The SLR and QSVM models were trained using two different training sets: 1) Monolayer Boundary (MB) set, which utilizes monolayer area, monolayer perimeter, and drug concentration as predictors 2) Discretized Window (DW) set, which utilizes endothelium area, endothelium perimeter, in each overlapping discretized windows and drug concentration as predictors.

## Results

### RMS Tractions are best predicted by the SLR model

Both ML models (SLR and QSVM) were generated using data from cells exposed to two different chalcone concentrations (0.2 μg/ml and 2 μg/ml). The values, coefficient of determination (R^2^), and root mean square error (RMSE) were calculated for different chalcone concentrations (0 μg/ml, 0.2 μg/ml, 1 μg/ml, and 2 μg/ml) based on the average from three unseen monolayers for each concentration. The results for 0 μg/ml and 1 μg/ml chalcone concentrations were broke down and shown separately in the model validation section comparing it with the corresponding actual experimental results.

#### Monolayer Boundary Predictor

A low R^2^ of 0.51 & 0.54 were seen for RMS tractions predicted by SLR and QSVM models for different chalcone concentrations (0 μg/ml, 0.2 μg/ml, 1 μg/ml, and 2 μg/ml). RMSE for the predictions from SLR and QSVM models were 6.5Pa and 5Pa for RMS tractions, respectively. The prediction accuracy of RMS tractions was the best for the QSVM model with R^2^ of 0.54 (from extended data table 1). The predicted RMS tractions distributions as a function of chalcone concentrations 0.2 μg/ml and 2 μg/ml is shown in figure 2 (a-g), (monolayer boundary training set - left panel). The corresponding averages were shown in extended data table 2 and figure 2g (monolayer boundary training column)

**Figure 1.**
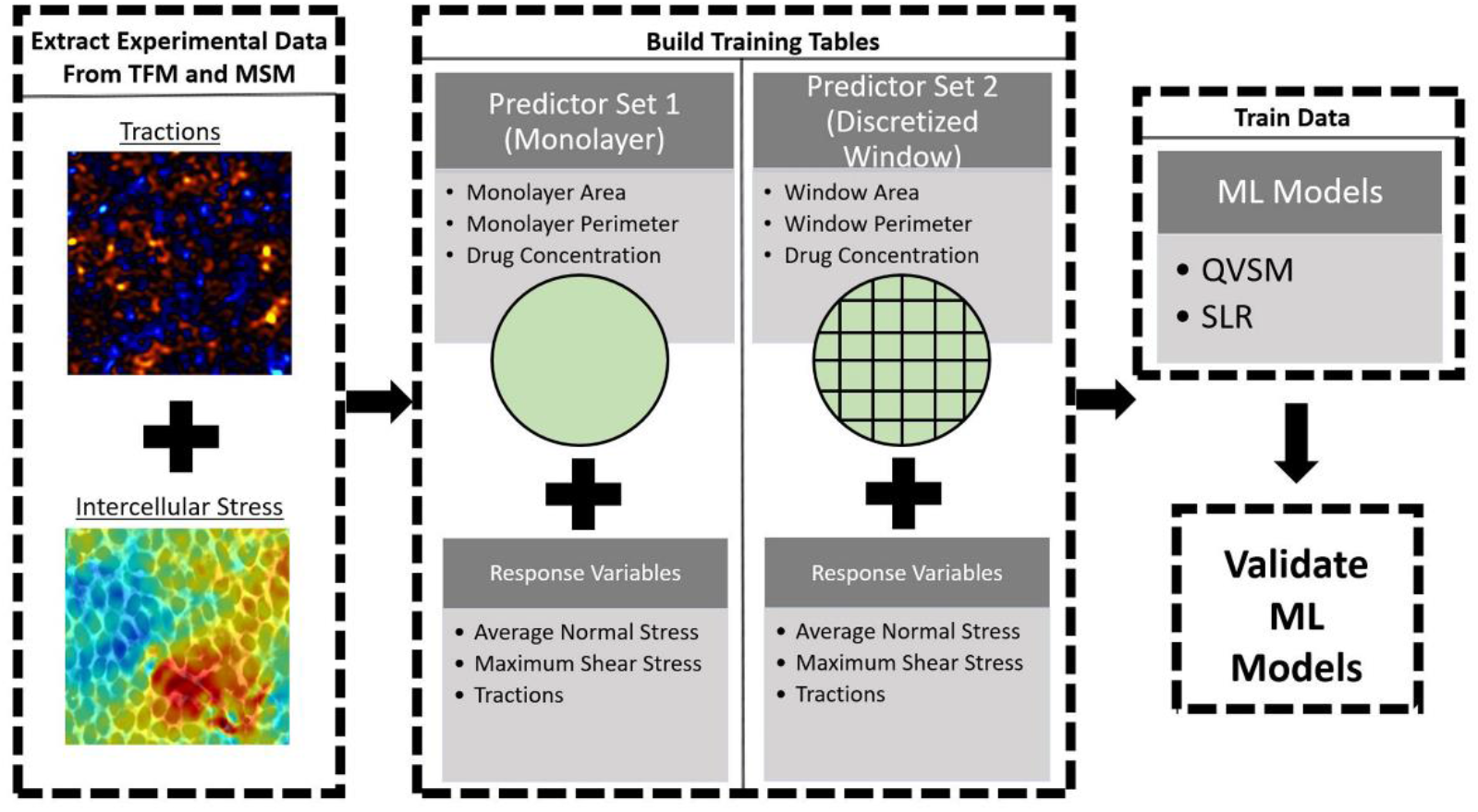
Flowchart showing implementation of machine learning approach to predict intercellular stress and tractions. The QSVM and SLR machine learning models were used along with two predictor sets: 1) Monolayer Boundary Set 2) Discretized Window Set.

**Figure 2.**
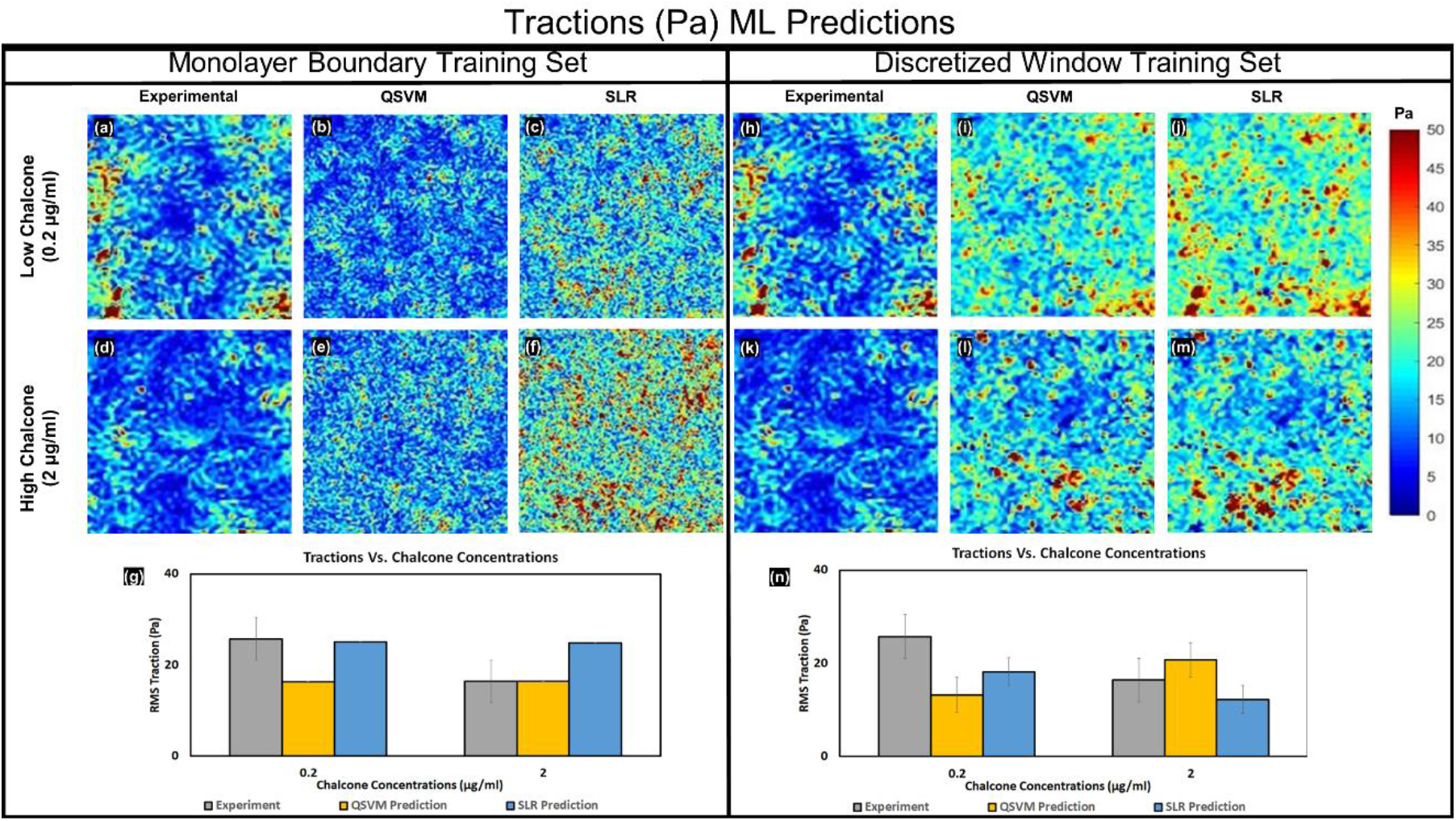
Predicted tractions using monolayer boundary training set and discretized window training set. Experimental RMS traction, QSVM, and SLR predicted RMS traction distributions for 0.2 μg/ml chalcone concentration (a-c) and 2 μg/ml chalcone concentration (d-f) and the corresponding averages from 3 samples for each condition with standard errors is shown in the bar plot (g) utilizing monolayer boundary training set. Experimental RMS traction, QSVM, and SLR predicted RMS traction distributions for 0.2 μg/ml chalcone concentration (h-j) and 2 μg/ml chalcone concentration (k-m) and the corresponding averages from 3 samples for each condition with standard errors is shown in the bar plot (n) utilizing discretized window training set.

#### Discretized Window Predictor

Similar to results discussed above, predictions were made using SLR and QSVM models for three unseen samples for 0.2 μg/ml and 2 μg/ml chalcone concentrations using discretized window training sets and compared against the experimental results (figure 2 (h-n), discretized window training set - right panel). The R^2^ values for RMS tractions predicted by SLR and QSVM were 0.85 and 0.008 for SLR and QSVM and 56Pa and 7.37Pa, respectively for different chalcone concentrations (0 μg/ml, 0.2 μg/ml, 1 μg/ml, and 2 μg/ml) as shown in extended data table 1. The combination of the SLR model along with the discretized window predictor gave us our best prediction of RMS tractions. The average tractions for 0.2 μg/ml and 2 μg/ml were shown in figure 2n and extended data table 2 in the discretized window sections respectively.

### Intercellular stresses are best predicted by QSVM model

#### Monolayer Boundary Predictor

The overall coefficient of determination (R^2^) for the average normal stress predicted by SLR and QSVM models for different chalcone concentrations (0 μg/ml, 0.2 μg/ml, 1 μg/ml, and 2 μg/ml) were 0.74 & 0.7 respectively. R^2^ for maximum shear stress predicted by SLR and QSVM were 0.88 & 0.84 respectively (extended data table 3, monolayer boundary training column). RMSE for the predictions from SLR and QSVM models were 24.18Pa and 27.5Pa respectively for average normal stress and 47.26Pa and 53.57Pa for maximum shear stress, respectively. Overall, the prediction accuracy was the best utilizing SLR model, for average normal and maximum shear intercellular stresses (R^2^ of 0.74 and 0.88) from extended data table 3, monolayer boundary training column. The predicted intercellular stresses (average normal and maximum shear) distributions for 0.2 μg/ml and 2 μg/ml chalcone concentrations were shown in figure 3 (average normal stress, a-c and g-i) and figure 3 (maximum shear stress, d-f and j-l) in the left panel (monolayer boundary training set) and the corresponding averages were shown in figure 3y (bar plots) and extended data table 4 in the monolayer boundary training sections.

**Figure 3.**
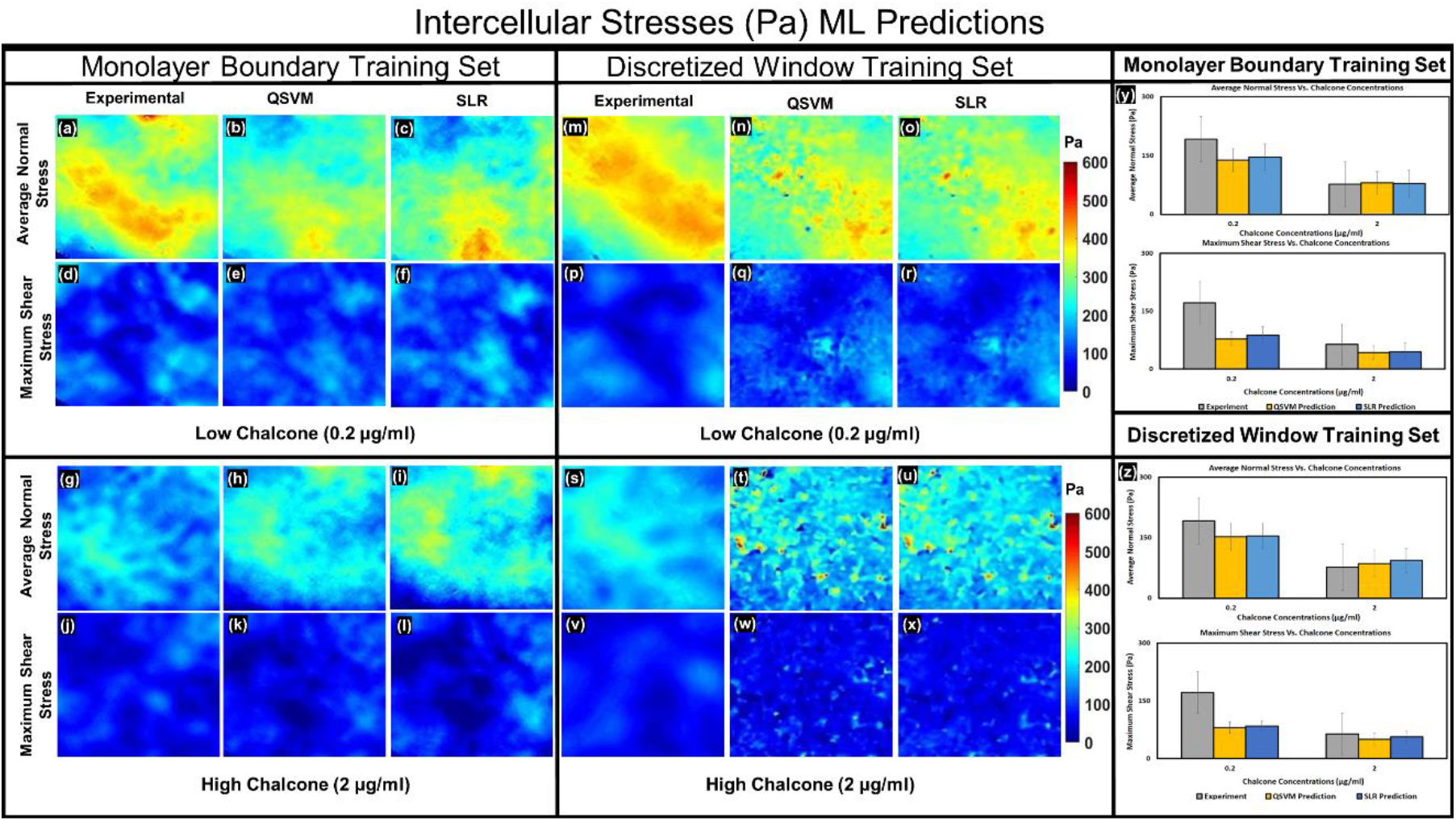
Predicted intercellular stresses using monolayer boundary training set and discretized window training set. Experimental average normal stress, QSVM, and SLR predicted average normal stress distributions for 0.2 μg/ml chalcone concentration (a-c) using monolayer boundary sets and (m-o) using discretized window training sets. Experimental maximum shear stress, QSVM, and SLR predicted maximum shear stress distributions for 0.2 μg/ml chalcone concentration (d-f) using monolayer boundary sets and (p-r) using discretized window training sets. Experimental average normal stress, QSVM, and SLR predicted average normal stress distributions for 2 μg/ml chalcone concentration (g-i) using monolayer boundary sets and (s-u) using discretized window training sets. Experimental maximum shear stress, QSVM, and SLR predicted maximum shear stress distributions for 2 μg/ml chalcone concentration (j-l) using monolayer boundary sets and (v-x) using discretized window training sets. Corresponding averages of the distributions from 3 monolayers for each condition with standard errors were shown in figure 3 y using monolayer boundary training sets and 3 z using discretized window training sets.

#### Discretized Window Predictor

The R^2^ values for average normal stress predicted by SLR and QSVM were 0.67 and 0.81, respectively. RMSE values for the maximum shear stress for SLR and QSVM were 24.49 Pa and 22.20Pa, respectively. Higher R^2^ values of 0.87 and 0.93 were seen for maximum shear stress predicted by SLR and QSVM models with RMSE of 49.06Pa and 52.67Pa, representing our best predicted values for different chalcone concentrations 0 μg/ml, 0.2 μg/ml, 1 μg/ml, and 2 μg/ml respectively. The predicted intercellular stresses: average normal and maximum shear distributions for 0.2 μg/ml and 2 μg/ml chalcone concentrations were shown in figure 3 (average normal stress, m-o and s-u) and figure 3 (maximum shear stress, p-r and v-x). The average values for the same were shown in extended data table 4 and figure 3z (bar plots) in the discretized window training sections.

### Validation of our ML model

We determined the prediction accuracy was to be best determined overall utilizing QSVM model for intercellular stresses (R^2^ of 0.81 and 0.93, respectively) and SLR model for RMS tractions (R^2^ of 0.85) along with the discretized window area predictor (extended data table 3 and table 1). Therefore, it was our next objective to evaluate the predictive capability of our ML model. To this end, we predicted intercellular stresses and rms tractions in response to 0 μg/ml and 1 μg/ml of chalcone and compared our predictions to experimental values obtained from these concentrations. We highlight the fact that 0 μg/ml and 1 μg/ml chalcone data was not used to train our model and the results we present here represent a true prediction and not a fit. Predicted spatial distributions of intercellular stresses and tractions of monolayers exposed to 0 μg/ml and 1 μg/ml compared to experimental results is shown in the figure 4 (monolayer boundary set, a-g) and figure 4 (discretized window set h-n) for RMS traction predictions. Intercellular stress predictions (average normal and maximum shear) were shown in figure 5 (monolayer boundary set, a-l and y) and figure 5 (discretized window set, m-x and z). Values obtained were displayed in extended data table 5 and table 6.

**Figure 4.**
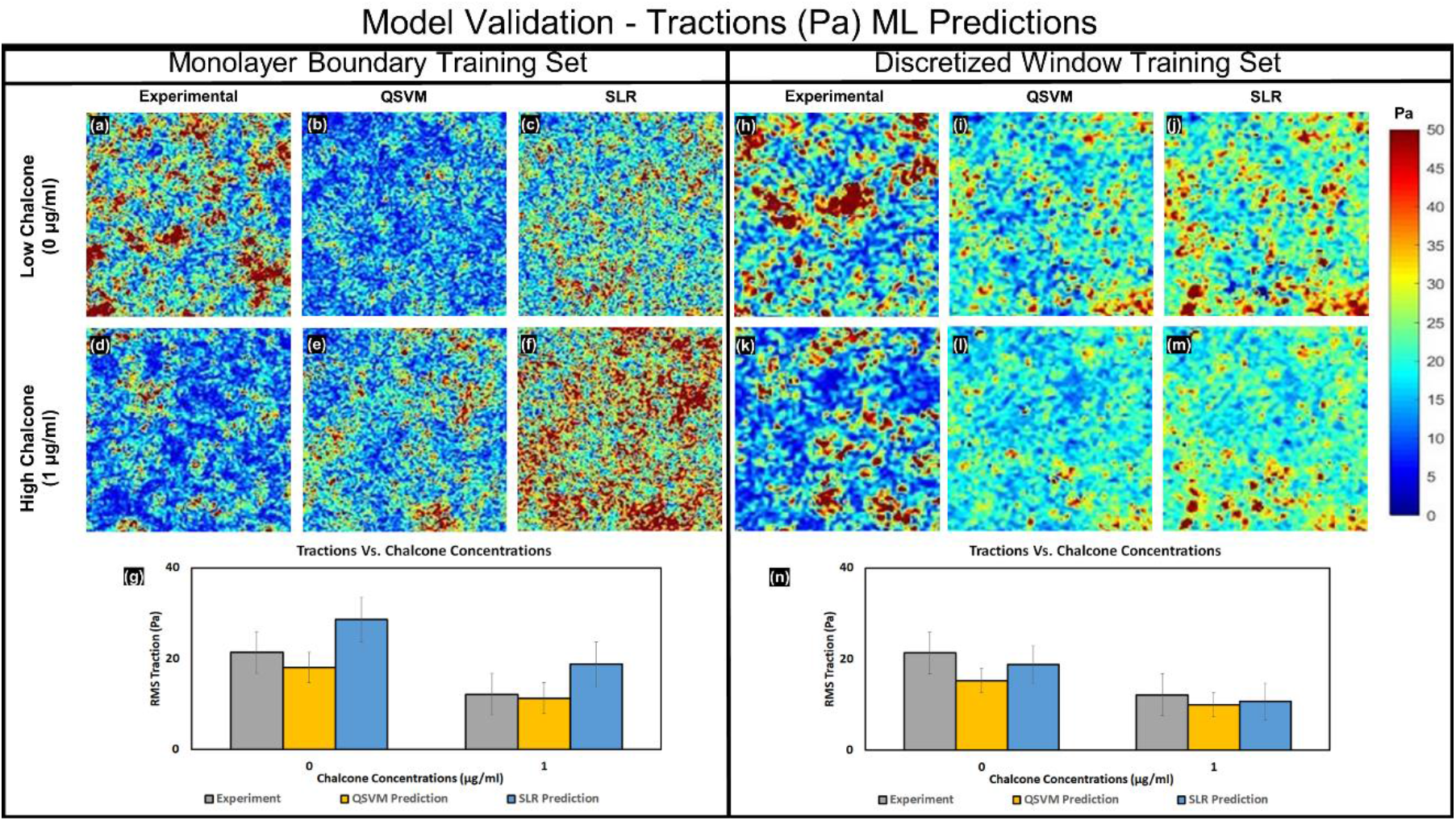
Model validation of tractions using monolayer boundary training set and discretized window training set. Experimental RMS traction, QSVM, and SLR predicted RMS traction distributions for 0 μg/ml chalcone concentration (a-c) and 1 μg/ml chalcone concentration (d-f) and the corresponding averages from 3 samples for each condition with standard errors is shown in the bar plot (g) utilizing monolayer boundary training set. Experimental RMS traction, QSVM, and SLR predicted RMS traction distributions for 0 μg/ml chalcone concentration (h-j) and 1 μg/ml chalcone concentration (k-m) and the corresponding averages from 3 samples for each condition with standard errors is shown in the bar plot (n) utilizing discretized window training set.

**Figure 5:**
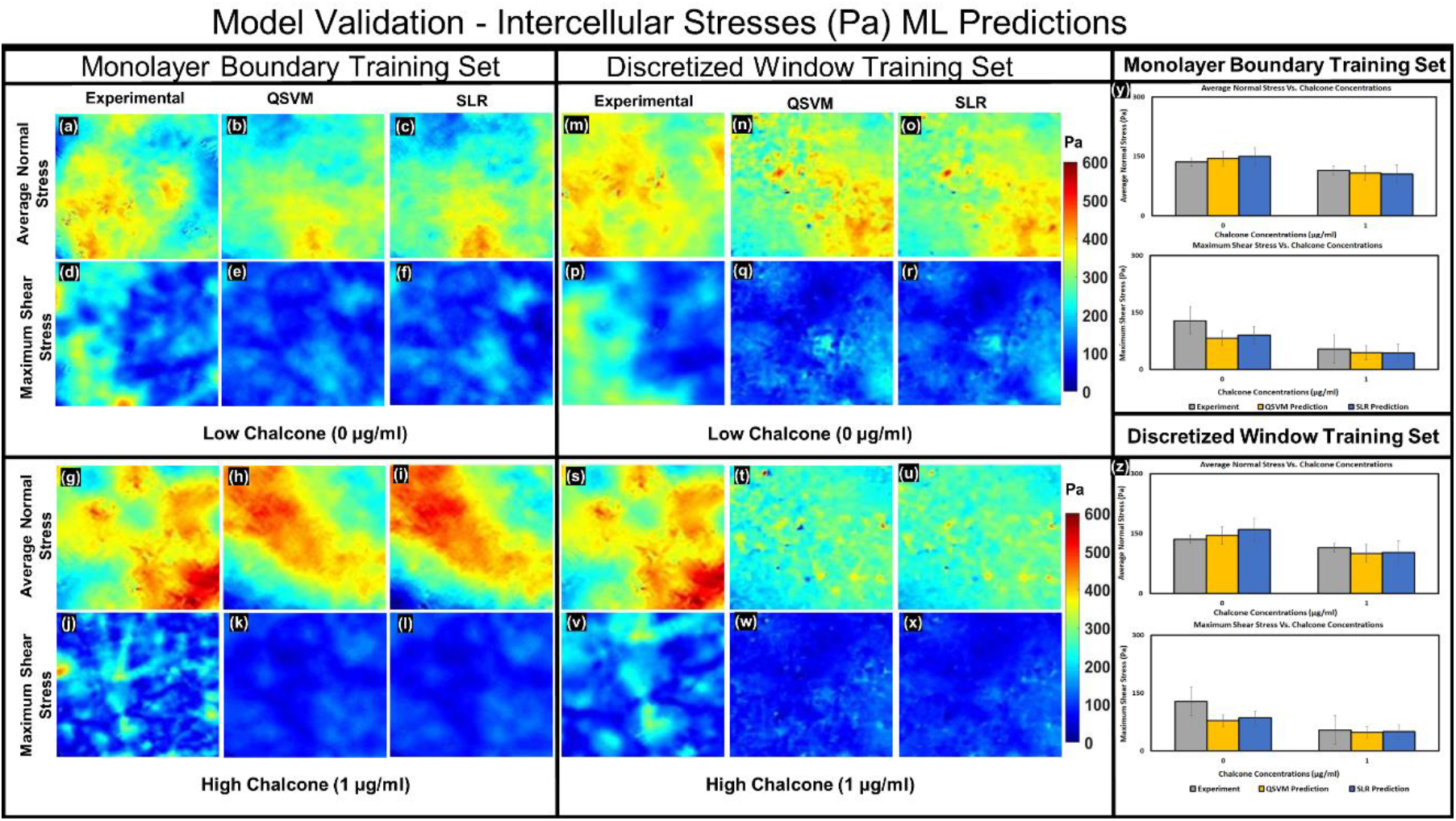
Model validation of intercellular stresses using monolayer boundary training set and discretized window training set. Experimental average normal stress, QSVM, and SLR predicted average normal stress distributions for 0 μg/ml chalcone concentration (a-c) using monolayer boundary sets and (m-o) using discretized window training sets. Experimental maximum shear stress, QSVM, and SLR predicted maximum shear stress distributions for 0 μg/ml chalcone concentration (d-f) using monolayer boundary sets and (p-r) using discretized window training sets. Experimental average normal stress, QSVM, and SLR predicted average normal stress distributions for 1 μg/ml chalcone concentration (g-i) using monolayer boundary sets and (s-u) using discretized window training sets. Experimental maximum shear stress, QSVM, and SLR predicted maximum shear stress distributions for 1 μg/ml chalcone concentration (j-l) using monolayer boundary sets and (v-x) using discretized window training sets. Corresponding averages of the distributions from 3 monolayers for each condition with standard errors were shown in figure 5 y using monolayer boundary training sets and 5 z using discretized window training sets.

## Discussion

In this paper, we present here both tractions and intercellular stresses as a function of monolayer morphology and drug concentration for the first-time using regression machine learning models QSVM and SLR with two different training sets, utilizing one of the monolayer boundary or discretized window sets. Overall, SLR models exhibited optimal predictive capability with monolayer boundary training sets for intercellular stress predictions while QSVM model did perform more optimally than SLR for discretized window training sets for the same. The R^2^ of intercellular stress predictions was the highest for QSVM model utilizing discretized window training set (R^2^ = 0.81 for average normal stress and R^2^ = 0.93 for maximum shear stress). SLR model utilizing discretized window training set had the highest R^2^ value of 0.85 for traction predictions. While on the one hand QVSM best predicted intercellular stresses on the other hand QVSM also predicted tractions the most poorly, yielding a R^2^ value of 0.008 for RMS traction predictions using QSVM. We believe this could be attributed to the fact that the discretized window training predictor was used and potential boundary effects from edges which were used to train the regression models. Discretized window training requires preprocessing to extract input variables from the new data (image) for making predictions while monolayer boundary method requires very little preprocessing for extracting the independent variables. The prediction time for a new (unseen) image was relatively same across the all the ML models built on the two different training sets (~10-20 min).

Out of the two proposed training sets, ML models built on discretized window training were very much sensitive to the outliers. In discretized window method, the predictions from all the three regression models (SLR, LSVM, and QSVM) were subject to a filter that excludes the outliers < −1000 Pa & > +1000 Pa (values corresponding to 0.15^th^, 99.95^th^ percentile) for intercellular stresses and > +600 Pa (values corresponding to the 99.95^th^ percentile) for RMS tractions to filter noise due to overfitting. Also, the most important feature of the discretized window method is the ability to generate time series predictions of tractions and intercellular stresses based on the changes in area & perimeter observed in each small overlapping grid across the entire field of view. The R^2^ especially for tractions can be improved by using more predictors such as substrate stiffness, cell orientation, cell velocity, etc. Also, training the models using a cropped monolayer section inside the circular monolayer samples could reduce the noise due to boundary effects and improve prediction accuracy. Time series prediction is possible with discretized window training but R^2^ was too low because of overfitting due to noise for both tractions and intercellular stresses. One of the future works is to reduce the boundary effects of the training sets by using cropped subsets inside the monolayer and employing cell velocity as an independent variable for time series predictions. However, including cell velocity as a predictor can increase the pre-processing and computation time compared to window-based and monolayer area predictors. Other future works include making time series predictions for expanding monolayer using monolayer boundary training sets and explore with more predictors such as cell velocity, cell orientation, curvature, cell area, ECM concentration, and substrate stiffness.

### Comparison of Predicted Results to Experimental Results

Predictions made for each response variable were computed for a new image with different monolayer area and drug concentration. The models were built on data from 4 monolayer samples with just two chalcone concentrations (0.2 & 2μg/ml). Predictions were made for new phase images with chalcone concentrations set to 0 & 1μg/ml respectively. Master average of the predicted results (RMS Traction, Average Normal Stress, Maximum Shear Stress) were compared with the actual master average of the results computed from TFM & MSM. Master average of the results is the average of the response variable in all the overlapping blocks that make up the image and the average taken across the entire image time series that follow in the MSM experiment. Predictions were made for 3 monolayer samples for each of the four chalcone concentrations, using the proposed ML models based on two different training sets and compared against the actual experimental result averages from 3 samples for each chalcone concentration obtained from MSM. Pearson correlation coefficient (R) was calculated in Excel 2016, using the formula: R = Covariance (A, B) / (Std. Dev A * Std. Dev. B). If the trend between MSM and predicted results were close, then we get high R values and vice versa. R^2^ (coefficient of determination) is a much-preferred metric in statistics which represents the % of variability in data that can be explained by the ML model. Root Mean Square Error (RMSE) was also computed for the predicted averages of the response variable against the experimental MSM data based on averages from 3 samples, across 60,516 grid points in each of the 292 images in the time series for each chalcone concentration.

## Conclusion

Intercellular stresses and tractions measured from a HUVEC monolayer can depend on a wide variety of factors such as monolayer area, substrate stiffness, cell area, curvature, external forces & biochemical substances, or drugs. Cell tractions have been correlated to factors like substrate stiffness, cell area, local curvature, and cell geometry [9]. Bayesian inversion stress microscopy (BISM) and Kalman inversion stress microscopy (KISM) were recently introduced to predict intercellular stresses from traction force microscopy. However, we believe this is the very first work to predict both tractions and intercellular forces from the independent variables: drug concentration, monolayer area/perimeter, and discretized window area/perimeter based on prior knowledge of tractions and intercellular stresses. We predicted both tractions and intercellular stresses using QSVM and SLR regression learners built on two different training sets. The coefficient of determination (R^2^) value was the highest for maximum shear stress compared to average normal stress and RMS tractions which was relatively consistent across different ML models based on averages from 3 unseen samples for each drug concentration. Inclusion of more predictors such as substrate stiffness should increase the prediction accuracy (R^2^) of the regression learners. With these promising results, we believe more accurate predictions can be obtained by adding additional predictors such as cell velocity, substrate stiffness, ECM concentration, for example, to improve the accuracy of the current models to make more reliable predictions of stress distributions and time series predictions instead of just the average trend. This work will be helpful for accelerating research in experimental drugs that target cell mechanical activity such as cellular contractility and tissue barrier strength and function. Proposed ML models could be applicable for testing cell mechanics of any anchorage dependent cells as a function of pharmacological and other morphological parameters that can influence cell mechanics.

## Materials and Methods

### Cell Culture

Human Umbilical Vein Endothelial Cells (HUVECs) were purchased commercially and cultured in Medium 200 supplemented with 1% penicillin-streptomycin (Corning) and Large Vessel Endothelial Supplement (LVES). HUVECS, Medium 200, and LVES were purchased from ThermoFisher. HUVECs were cultured on 0.1% Gelatin (Sigma-Aldrich) coated flasks at 37° C and 5% CO_2_.

### Preparation of Polyacrylamide Gel and Cellular Micropatterning

The protocol for preparing Polyacrylamide (PA) gels can be found in Steward et al. [37]. Glass bottom petri dishes (35 mm, Cellvis) were treated with bind silane solution for 45 minutes after which the dishes were rinsed with deionized water and air-dried. PA solution is made by mixing ultra-pure water, 40% acrylamide (Bio-Rad), 2% bis-acrylamide (Bio-Rad), and fluorescent beads (yellow or Texas red with ~0.5 μm diameter, Invitrogen). The stiffness of the PA gel can be fine-tuned by changing the ratio of Bis(acrylamide) and acrylamide solutions. The PA solution is kept in a vacuum chamber for 40 minutes. 10% ammonia persulfate and TEMED (N,N,N’,N’-tetramethylethane-1,2-diamine) were added to the de-gassed PA solution which initiates polymerization reaction. After the addition of ammonia persulfate, TEMED the PA solution is mixed well and plated on the petri dish wells. Hydrophobic coverslips were then placed on top of the PA solution after which the dishes were inverted to allow more fluorescent beads to settle on top layer of the polymerizing gel. The subsequent PA gels with stiffness ~1.2 kPa and height ~100 μm were used for the experiments as described by Stroka et al. [38].

### Cellular Micropattern Preparation

Polydimethylsiloxane (PDMS) was used to fabricate thin micropatterns as described previously in [37]. A thin cross-section of PDMS (Dow Corning) was prepared by mixing silicone base with a curing agent (20:1) and the mixture was then poured into a 100 mm petri dish. The PDMS mixture in the petri dish with no air bubbles was then incubated at 70° C overnight. Thin, circular cross-sections of cured PDMS (16 mm) were fabricated using a hole puncher. Small 2 mm holes were made on the circular PDMS section using a biopsy punch. The fabricated micropatterns were gently placed on the top layer of the PA gels.

### SANPAH Burning & Col I-FN Treatment

The petri dish samples with PDMS micropatterns stamped on PA gels were then subject to treatment with sulfosuccinimidyl-6-(4-azido-2-nitrophenylamino) hexanoate (Sulfo-SANPAH; Proteochem) dissolved in 0.1 M HEPES buffer solution (Fisher Scientific) and kept under a UV lamp for 8 mins. After SANPAH and UV treatments, the samples were treated with type Collage I (Col-I).

After the treatment of Col-I (Advanced Biomatrix) overnight at 4° C, the excess protein solution was carefully removed and HUVECs were seeded at a density of ~50 × 10^4^ cells/mL. After 60-75 mins, micropatterns were cautiously removed using a tweezer. The HUVEC’s monolayer samples were incubated at 37° C and 5% CO_2_ for at least 24 hours to allow enough time for the formation of cell-cell junctions prior to experimentation.

### Pharmacological Perturbation Experiments

We have previously demonstrated tractions and intercellular stresses to exhibit a dose-dependent response to 2,5-dihydroxychalcone (chalcone) [39]. Chalcone is a unique drug in that it has been reported to solely disrupt the gap junction connexin 43 [39]. Therefore, for this study HUVECs were seeded at a density of 50 × 10^4^ cells/mL onto polyacrylamide gels for at least 36. After this time, independent experiments were conducted where chalcone was added at the following concentrations: 0.2 μg/mL (low concentration) and 2 μg/mL (high concentration). Experiments performed to validate our machine learning model consisted of controls and exposing HUVECs to 1 μg/mL of chalcone.

### Time Lapse Microscopy

Time lapse microscopy was performed using a Zeiss inverted microscope with a 10X objective and Hamamatsu camera. Fluorescent and phase contrast images were acquired at 5-minute intervals for 1 hour prior to the addition of chalcone. After this time, medium with chalcone was added at the concentrations mentioned above and imaging was done for an additional 5 hours at 5-minute intervals after removing the control medium. These experiments culminated with HUVEC monolayers being treated with 10X Trypsin to detach the cells from the substrate and acquire a “stress free” image of our gel surface, an image essential for traction calculation.

### Traction Force Microscopy (TFM) & Monolayer Stress Microscopy (MSM)

First, in-plane displacement of fluorescent beads located on the top surface of the gel was computed using a custom-written Particle Image Velocimetry (PIV) routine. A window size 32 pixels with an overlap of 0.75 was utilized for each Region of Interest (ROI). Cross correlation between each window in the reference image with no cells attached to gel (stress-free configuration) was computed against a window occupying the same coordinates in the fluorescent image with cells attached (stressed configuration) sequentially, across all the window blocks. The displacements were calculated in the x and y coordinate system in pixels using an iterative cross-correlation function. The displacement calculated from peak cross-correlation function between each reference-fluorescence window pair was assigned to the center coordinates of those windows [3]. Traction force Microscopy [40] and monolayer stress microscopy [7, 18] were used as described by Butler et al. and Trepat et al. to calculate the cell-substrate tractions and cell-cell intercellular stresses, respectively. Briefly, the gel deformations described above were used to calculate the tractions and the intercellular stresses were subsequently recovered from traction force maps by using straightforward force balance equations imposed by Newton’s law [19]. We computed the local two-dimensional stress tensor within the monolayer by converting the maximum principal stress (σ_max_) and minimum principal stress (σ_min_) along the principal plane by rotating the local coordinate system along the principal orientation. The average normal stress (σ_max_ + σ_min_)/2 and maximum shear stress (σ_max_ - σ_min_)/2 were calculated at each point in the elements within the monolayer.

### Building of Training Tables

Experimental TFM and MSM data used for building our training tables were solely obtained from the 0.2 μg/mL chalcone and 2 μg/mL chalcone experiments. We used results gathered from four-time lapse experiments (two per each chalcone concentrations: 0.2 μg/mL and 2 μg/mL chalcone experiments respectively) and concatenated the data sets row-wise into one large data set. The data gathered from TFM and MSM experiments were in the form of a cropped, 246×246 square matrix, which contained approximately 60516 data points per time series once converted into column form. This data column was subsequently used to build training tables for our predictor and response variables. The three response variables were RMS tractions, maximum principal stress, and minimum principal stress. Column training tables were created separately for each of these response variables with the help of predictors before training them with ML models. Predictors utilized were drug concentration and either of the two morphological predictors: monolayer boundary, or discretized monolayer windows. In addition, the Principal Component Analysis (PCA) tool in MATLAB was used to confirm that our selected predictors explained more than 95% of variability in the data.

### Monolayer Boundary Training Table

The monolayer boundary training table was generated from the monolayer area. A contour of the monolayer shape was obtained from phase contrast images and converted to a binary image. Utilizing this binary image monolayer area & perimeter were then calculated using the ‘regionprops’ command in MATLAB. The monolayer area & perimeter remain the same for all the time frames as our monolayer was seeded as a constrained, micropattern during experiments. Monolayer boundary data cannot capture individual HUVEC movement and individual HUVEC geometry within the monolayer since measurements are only made for the entire monolayer as a whole.

Furthermore, single monolayer area and perimeter does not change the monolayer boundary data set can only train ML models to predict the average values of our response variables. Ramifications of this include the lack of ability to predict dynamic behavior of our response variables, which would be revealed through a time series. To overcome the monolayer boundary data’s lack of ability to predict time series information we developed the discretized window training table.

### Discretized Window Training Set

Discretized Window training data was generated by converting binarized phase images of cellular monolayers into multiple overlapping grids which we call here “windows”. For any given frame we used a grid resolution of 32 × 32 to generate our windows and from these windows we calculated an area and perimeter, which was then stored as a column matrix for each image. The area and perimeter of each discretized window gives information about coverage of endothelium within that window. We choose this method as computing cell area and perimeter in an automated fashion would be difficult and time consuming as these cell properties change from frame to frame. The discretized window training method eliminates the need to track individual cell properties and is 2.6x faster compared to our traditionally used FFT (Fast Fourier Transforms)-based, cross-correlation method used for cell displacement tracking for a single image pair of 32 sq. pixels each (supplementary figure 4).

### Selection of ML Models

The generated predictor variables and response variables mentioned above were utilized into several machine learning models in MATLAB using the Machine Learning toolbox. Since the data was relatively small, we were able to utilize all the available ML models in MATLAB. A five-fold variable cross validation was chosen to validate the models where the inputted training table was randomly divided into five groups (refer supplementary section 3) and one group was held as the “test data” from which the model makes predictions after training data from the remaining four sets. ML models that 1) had the highest R^2^ and lowest RMSE values when predicting our response variables (tractions and intercellular stresses) and 2) were most sensitive to our predictor variables (area, perimeter, and drug concentration) served as our selection criteria. Utilizing this criteria, the Support Vector Machine (SVM) and Stepwise Linear Regression (SLR) based models emerged as our best candidates for predicting TFM and MSM data as a function of area, perimeter, and drug concentration. Trained models based on SVM and SLR were found to be sensitive to new data with less overfit and high R^2^ values compared to the other models. SVM and SLR are described below.

### Support Vector Machine

Support Vector Machine (SVM) is widely used in classification and regression analysis. SVM makes predictions by utilizing various kernelization techniques to effectively find the hyperplane that separates the support vectors by transforming the data in higher dimensions in classification problems. In regression, the hyperplane is the best fit line or curve that effectively fits most of the points or support vectors. SVM can use different kernels such as linear, quadratic or gaussian to compute the transformation in higher dimensions at a reduced computational cost, where it is easier to find the hyperplane that is closer to most of the data points. SVM optimizes the hyperplane so that the distance from each support vector is minimized within the chosen decision boundary. The decision boundary encompasses the data that are closer to the hyperplane. A margin of tolerance (ε) can be inputted by the user to increase the tolerance level from the decision boundary. SVM is well-equipped to handle complex, non-linear data, robust to outliers and has better prediction accuracy compared to linear regression models. However, SVM is susceptible to noise and not preferred for very large or very small data sets [41].

### Stepwise Linear Regression

Stepwise Linear Regression (SLR) evaluates the independent or exploratory variables one by one through a forward selection rule (variables added at each step), backward-elimination rule (all variables included), or a bi-directional rule (combination of both forward selection and backward elimination) by computing the t-statistics for the coefficients of the selected variable at each step. SLR regression model helps choose the best independent variables efficiently based on statistical significance. Although very effective in minimizing the number of predictors, SLR model is prone to choosing wrong variables if there is a large number of predictors and small amount of data [42].

## Author Contributions

M.M.I ran experiments and provided data for building the machine learning model. V.A.S.B. built the machine learning models, analyzed data and generated figures. V.A.S.B and R.L.S. interpreted the data and V.A.S.B., M.M.I. and R.L.S. contributed to the writing of the manuscript.

## Data and Code Availability

The data and code that support the findings of this study are available from the corresponding author, [R.L.S], upon reasonable request.

## Acknowledgements

This work was funded by the National Heart, Lung, And Blood Institute of the National Institute of Health under award K25HL132098. This material is based on work supported by the National Science Foundation CAREER Award under Grant No. 2045750.

## Figures and Tables

**Extended Data Table 1.** R^2^ and RMSE of RMS Traction predictions using ML models from monolayer boundary and discretized window training sets for different chalcone concentrations (0, 0.2, 1, and 2 μg/mL) based on average from 3 monolayer samples for each condition.

| Monolayer Boundary Training |  |  | Discretized Window Training |  |
| --- | --- | --- | --- | --- |
| RMS Traction | SLR Model | QSVM Model | SLR Model | QSVM Model |
| $R^2$ | 0.51 | 0.54 | 0.85 | 0.008 |
| RSME (Pa) | 6.5 | 5.0 | 4.56 | 7.37 |

**Extended Data Table 2.** RMS Traction predictions using ML models from monolayer boundary and discretized window training sets for 0.2 and 2 μg/mL chalcone concentrations based on average from 3 monolayer samples for each condition

| Experimental Results |  | Monolayer Boundary Training |  | Discretized Window Training |  |
| --- | --- | --- | --- | --- | --- |
|  |  | SLR Model (Pa) | QSVM Model (Pa) | SLR Model (Pa) | QSVM Model (Pa) |
| 0.2µg/ml | 25.7±5 | 25±5 | 16.3±2 | 18.2±1 | 13.2±1 |
| 2 µg/ml | 16.4±4 | 24.8±7 | 16.4±3 | 12.2±1 | 20.7±5 |

**Extended Data Table 3.** R^2^ of Intercellular stress and RMS Traction predictions using ML models from monolayer boundary and discretized window training sets for different chalcone concentrations (0, 0.2, 1, and 2 μg/mL) based on average from 3 monolayer samples for each condition.

| Monolayer Boundary Training |  |  | Discretized Window Training |  |
| --- | --- | --- | --- | --- |
| Avg Normal | SLR Model | QSVM Model | SLR Model | QSVM Model |
| $R^2$ | 0.74 | 0.7 | 0.67 | 0.81 |
| RSME (Pa) | 24.18 | 27.5 | 24.49 | 22.20 |
| Max Shear | SLR Model | QSVM Model | SLR Model | QSVM Model |
| $R^2$ | 0.88 | 0.84 | 0.87 | 0.93 |
| RSME (Pa) | 47.26 | 53.57 | 49.06 | 52.67 |

**Extended Data Table 4.** Intercellular stress predictions using ML models from monolayer boundary and discretized window training sets for 0.2 and 2 μg/mL chalcone concentrations based on average from 3 monolayer samples for each condition

| Experimental Results |  | Monolayer Boundary Training |  | Discretized Window Training |  |
| --- | --- | --- | --- | --- | --- |
| Average Normal Stress<br>(Pa) |  | SLR Model (Pa) | QSVM Model<br>(Pa) | SLR Model<br>(Pa) | QSVM Model<br>(Pa) |
| 0.2µg/ml | 192±32 | 146.6±8 | 138.2±8 | 154.4±7 | 152±7 |
| 2 µg/ml | 76.75±14 | 78.8±5 | 80.5±2 | 93.6±1 | 86.2±0.4 |
| Experimental Results |  | Monolayer Boundary Training Set<br>Predictions |  | Discretized Window Training<br>Set Predictions |  |
| Maximum Shear Stress<br>(Pa) |  | SLR Model (Pa) | QSVM Model<br>(Pa) | SLR Model<br>(Pa) | QSVM Model<br>(Pa) |
| 0.2µg/ml | 172±73 | 88.5±6 | 78.4±4 | 84.3±4 | 80.7±4 |
| 2 µg/ml | 63.8±4 | 44.6±2 | 44.6±1 | 56.6±3 | 50.6±2 |

**Extended Data Table 5.** Intercellular stress predictions using ML models from monolayer boundary and discretized window training sets for different chalcone treatment concentrations based on average from 3 monolayer samples for each condition

| **Experimental Results**<br>**Average Normal Stress (Pa)** |  | **Monolayer Boundary Training** |  | **Discretized Window Training** |  |
| --- | --- | --- | --- | --- | --- |
|  |  | **SLR Model (Pa)** | **QSVM Model (Pa)** | **SLR Model (Pa)** | **QSVM Model (Pa)** |
| **0µg/ml** | 136.19±25 | 150±13 | 144.6±9 | 160±7 | 145.3±7 |
| **1µg/ml** | 114.71±3 | 105.5±4 | 107.6±5 | 103.2±7 | 100.4±8 |
| **Experimental Results**<br>**Maximum Shear Stress (Pa)** |  | **Monolayer Boundary Training Set Predictions** |  | **Discretized Window Training Set Predictions** |  |
|  |  | **SLR Model (Pa)** | **QSVM Model (Pa)** | **SLR Model (Pa)** | **QSVM Model (Pa)** |
| **0µg/ml** | 129±20 | 90.7±6 | 82.3±5 | 85.9±4 | 78.6±1 |
| **1µg/ml** | 54.3±4 | 43.6±6 | 44.8±5 | 50.8±6 | 48.3±6 |

**Extended Data Table 6.** RMS Traction predictions using ML models from monolayer boundary and discretized window training sets for different chalcone treatment concentrations based on average from 3 monolayer samples for each condition

| Experimental Results<br>RMS Traction (Pa) |  | Monolayer Boundary Training |  | Discretized Window Training |  |
| --- | --- | --- | --- | --- | --- |
|  |  | SLR Model (Pa) | QSVM Model (Pa) | SLR Model (Pa) | QSVM Model (Pa) |
| 0µg/ml | 21.4±3 | 28.6±6 | 18±2 | 18.8±1 | 15.3±1 |
| 1µg/ml | 12.2±0.4 | 18.8±2 | 11.3±1 | 10.6±1 | 10±1 |

